# The York Gospels: a one thousand year biological palimpsest

**DOI:** 10.1101/146324

**Authors:** Matthew D. Teasdale, Sarah Fiddyment, Jiří Vnouček, Valeria Mattiangeli, Camilla Speller, Annelise Binois, Martin Carver, Catherine Dand, Timothy P. Newfield, Christopher C. Webb, Daniel G. Bradley, Matthew J. Collins

## Abstract

Medieval manuscripts, carefully curated and conserved, represent not only an irreplaceable documentary record but also a remarkable reservoir of biological information. Palaeographic and codicological investigation can often locate and date these documents with remarkable precision. The York Gospels (York Minster Ms. Add. 1) is one such codex, one of only a small collection of pre-conquest Gospel books to have survived the Reformation. By extending the non-invasive triboelectric (eraser-based) sampling technique eZooMS, to include the analysis of DNA we report a cost effective and simple-to-use biomolecular sampling technique. We apply this combined methodology to document for the first time a rich palimpsest of biological information contained within the York Gospels, which has accumulated over the 1,000 year lifespan of this cherished object that remains an active participant in the life of York Minster. This biological data provides insights into the decisions made in the selection of materials, the construction of the codex and the use history of the object.

## Introduction

Illuminated manuscripts are objects of great worth and value, often in the past elaborately decorated and bound, emphasising their importance, not only as literary texts but also as physical objects of intrinsic and spiritual value. Moreover, a contemporaneous collection of animal skins bound together provides a remarkable biological resource, which may inform upon the husbandry of the animals and in turn shed light on the assembly of the codex. The utility of parchment documents as a store of biological information is confirmed by a number of molecular studies, which have successfully retrieved DNA sequences from parchments and produced comparisons with modern reference populations of cattle, sheep and goat [1–4]. These analyses have utilised isolated parchment fragments, which are then digested as part of the DNA extraction process. Understandably, such studies have not yet included bifolio from bound volumes.

In the light of the vast potential afforded by ancient manuscripts several authors are experimenting with non-destructive sampling of documents. Whilst most of these methods involve some form of spectroscopic analysis, novel approaches which release molecules from the surface, such as the use of synthetic gel films [5,6] have been reported, and this work can be seen as part of a wider push to develop sampling methods for material culture [7–11]. These studies also form part of an increasingly sophisticated analysis of the conservation status of objects, which includes analyses of the microflora within buildings and upon objects. In the case of parchment, this includes concerns over both parchment deterioration and risks to personnel health caused by mould (*hyphomycetous* fungi) [6,12–14].

Analyses of the traces left by the handling and use of an object are common in Palaeolithic and Neolithic archaeology and have been applied to personal adornment artefacts [e.g. 15,16] as well as stone tools. Much less common in the historical period, they can illuminate the choices made in the selection of raw materials, and the life history of an object [17]. A pioneering application of this analysis is Kate Rudy’s [18] use of densitometry to map the discoloration of parchment caused by repeated usage. Her work highlights a third level of biological data, the grime on the surfaces of parchments that attests to handling. In some cases, folia have experienced much more active intervention, such as devotional kissing or rubbing, activities that are likely to leave distinct microbial traces, which can be explored using biomolecular methods.

The analysis of both the raw materials of the codex (*i.e*. the skins selected from flocks and herds), and the microorganisms on the object (which may highlight its use history and conservation risk) must be compatible with conventional conservation treatment. Our contribution to this field has been to develop [19] and here to refine a triboelectric (eraser-based) sampling method, to recover first protein and now DNA from parchment. Dry cleaning with PVC erasers is a common and widely used conservation technique [20], and we analyse the waste material from this process, which would otherwise be discarded.

We apply our approach to document for the first time the vast array of biological information contained with a single codex, recovering both DNA and proteins from the dry eraser waste of skins from the York Gospels (York Minster Ms. Add. 1). One of only a small collection of pre-conquest (1066 CE) Gospel books to have survived the Reformation, the York Gospels are thought to have been written around the turn of the first millennium in the scriptorium of St. Augustine's monastery Canterbury [21] and then brought to York by Archbishop Wulfstan *circa* 1020 CE. The text contains all four gospels of the New Testament; a letter from King Cnut (r.1016-1035 CE) and documents concerning land ownership. It is the only surviving English Gospel book to contain the oaths taken by the deans, archdeacons, canons and vicars choral, dated to the 14th-16th century and is still used in ecclesiastical ceremonies today. Our study presents the rich palimpsest of biological information housed within this document, which we then compare to six legal documents with different curatorial histories from two collections held at the Borthwick Archive (York, UK).

**Figure 1:**
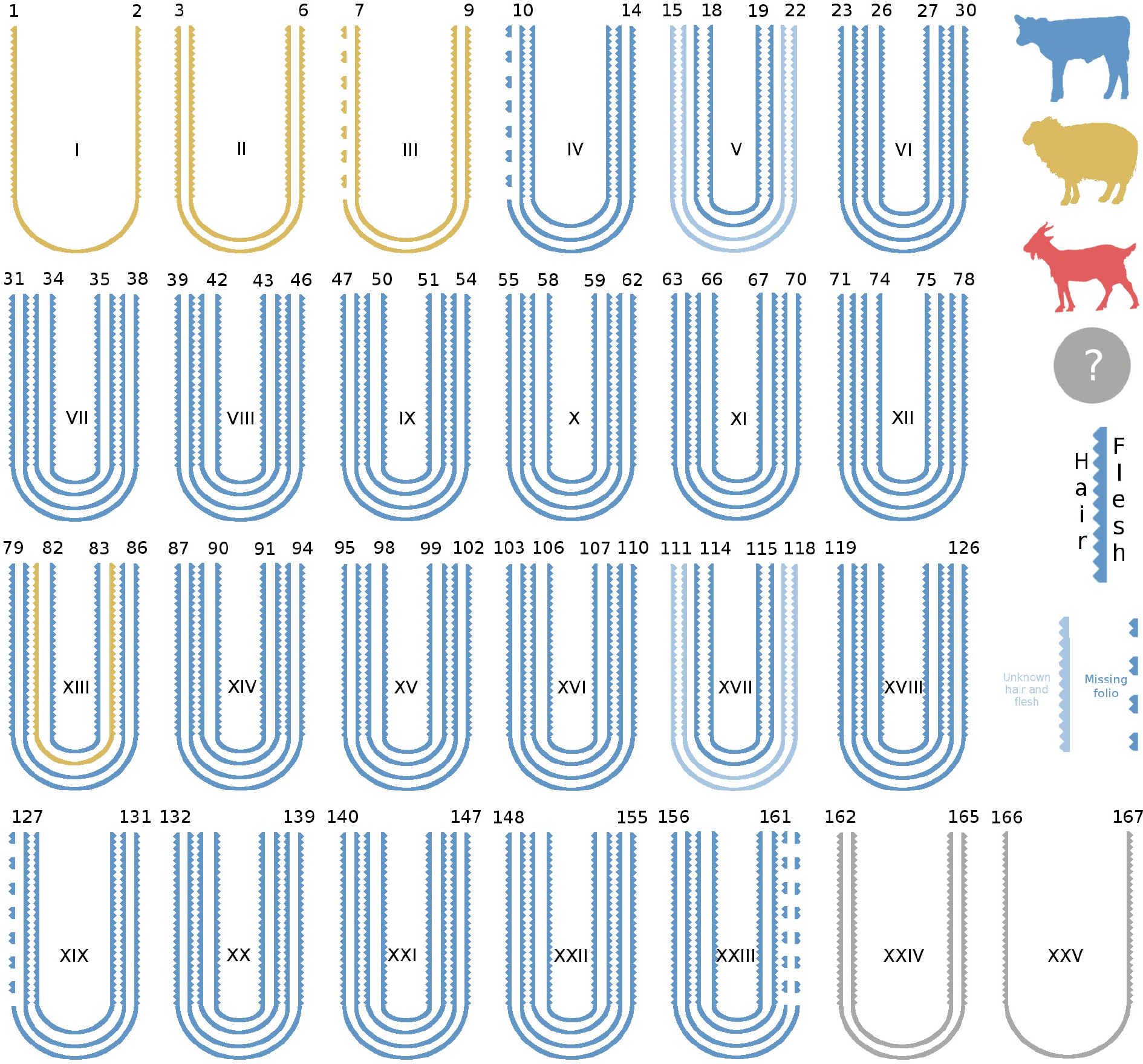
Structure of the quires comprising the manuscript and species identification of the bifolia using eZooMS.

## Results

### Species Identification

#### Protein Analysis

This is the first systematic application of eZooMS [19] to identify the source species of every bifolia contained within a single codex (Supplementary table 1). The York Gospels were found to be composed of two animal species: the original gospels are of calfskin (except for one bifolia made of sheep) and the later additions (C 14^th^) are made exclusively of sheepskin (figure 1, supplementary figure 1). Due to extensive conservation treatments carried out in the 1950s that involved covering entire folios in silk gauze, we were unable to determine the species for the final quire and flyleaves (folios 162-167).

#### DNA Analysis

DNA extraction was attempted from eight bifolio of the York Gospel from which large volumes of eraser waste (150-250ul) had been generated; sufficient DNA concentrations were recovered from all samples for successful library preparation and high-throughput sequencing. Initial species identification was undertaken with FastQ Screen and in all cases, the genetic species assignments agreed with those produced through eZooMS (supplementary table 2). One sample (folio 158, Quire XXIII), which had previously provided an inconclusive species identification of calf during the proteomic analysis was also assigned as calf in the genetic analysis. However, this page is one of those that suffered extensive invasive conservation, and even this combined assessment remains somewhat tentative. Insufficient data was recovered from two samples taken from the 14^th^C additions to the Gospels to confidently assign the source species using solely genetic methods, however both samples were previously conclusively identified as sheep via eZooMS.

### Genetic analysis

DNA sequences recovered from the parchment samples were stringently (supplementary table 3, supplementary methods) aligned to the genome of the identified host (production) species to estimate the proportion of endogenous DNA retained within the bifolio of the York Gospels (supplementary methods). This analysis resulted in a mean endogenous percentage of 19.3% (range 0.7-51.4%) over all the samples (supplementary table 3). For one bifolio, folio 125 (quire XVIII), 51.4% of reads could be aligned to the genome of the source species (cow), however this assignment fell to 5.6% when filters for read mapping quality were applied (supplementary table 3). While some loss of mapped reads is expected post filtering, this extreme reduction suggests that there may be a bias in DNA sequence preservation or retrieval from the manuscript; one that in this case favoured repetitive genomic regions over more gene rich euchromatic regions. This apparent taphonomic bias may be a reflection of the harsh alkaline treatment that is an integral part of the parchment production process, which might selectively degrade the more loosely packed euchromatin over the tightly bound heterochromatin (Supplementary figure 2).

A mapDamage2.0 [22] analysis was completed on the filtered host reads recovered from the York Gospels to explore DNA damage patterns and recovered DNA authenticity (supplementary figure 3). All samples were found to have characteristic markers of degraded ancient DNA with an increase in deamination at the end of reads. However, the frequency of these modifications is much lower than what would be predicted for bone of a similar age [23]. Given that the DNA fragment lengths are also short in these recovered sequences (supplementary figure 4), this result may suggest that different DNA degradation processes are active in parchment and bone.

Sufficient sequence data was recovered from three samples (Fol. 13, Fol. 101 and Fol. 125) to permit further population genetic analyses. These samples were placed onto a reference dataset of modern cattle [24] in a principal component analysis (supplementary figure 5a and b) using the LASER2.0 software [25,26]. Although, only a relatively small number of variants could be called in each of the three samples they can be seen to cluster with modern cattle breeds of Northern Europe (supplementary figure 5b). Specifically, the sample with the highest genomic coverage folio 101 (2,139 bovine HD snps called) falls just outside a cluster of Norwegian red and Holstein animals, with the two lower coverage samples folio 13 and folio 125 falling outside of the major European distribution (supplementary figure 5b). The position of folio 13 and 125 likely reflects the limited SNP recovery in these samples, however it could also be an indication of true genetic diversity in these animals. As further studies of geographically and temporally localised animal bone reveal the genetic landscape of medieval cattle, we will be in a stronger position to localise these and future parchment objects.

Sex identification [27] was attempted for all eight bifolio sampled for DNA analysis (supplementary table 2), with high confidence assignments being deduced for five. Four out of the five reliably typed animals in the original Gospel document (prior to the later *circa* 14^th^ C additions) were found to be female.

### Exogenous DNA

#### Human

As the novel non-invasive DNA sampling technique utilised in this study samples molecules from the surface of documents, the exogenous/environmental DNA residing on the York Gospels was explored. Firstly, to discern an estimate for the upper bounds of human DNA on the document, reads recovered from the Gospels were aligned to the human genome (hg19). The average recovery of human DNA sequences over the whole document was 11.1% (range 4.2 - 20.8%) (supplementary table 4), almost twice that of six control samples-legal documents (title deeds) held in the Borthwick Archive-sampled using identical methods (average 5.6%, range 1.9-12.8%). This lower proportion of human DNA residing on the control samples may reflect the use history of the documents, and the less intensive handling of legal deeds compared to the York Gospels. Sequences that aligned solely to the human genome show a reduction in DNA damage patterns compared to those from the host animal skin itself (supplementary figure 3), potentially indicating their more recent origin from ongoing handling. However, the overall DNA fragment lengths of the endogenous and exogenous molecules are comparable, with a possible slight shift to lower fragment lengths in the endogenous (parchment) molecules (supplementary figure 4).

For two York Gospel samples (Fol. 6 and Fol. 158) over 19% of recovered reads were mapped to the human genome (supplementary table 5), this dropped to 16% when filters for mapping quality were applied (supplementary table 5). The reduction in alignment percentage due to mapping quality filtering is not as dramatic as that seen in the endogenous (parchment) DNA, a result consistent with the idea that these molecules were deposited after parchment manufacture. Folio 158 (15.3%) was the subject of extensive and invasive conservation treatment in the last century and the exogenous DNA fragment lengths from this parchment show striking periodicity consistent with histone wrapping, [28,29] a result that may be indicative of a significant amount of DNA degradation (supplementary figure 4). Folio 6 (17.5%) contained the "Oaths of a subdean in person, a canon and prebendary in person, and an archdeacon", passages which we would anticipate to be more regularly read/handled than the other folia studied.

#### Microbiome

To further classify the York Gospels metagenome, recovered sequences were analysed using two independent metagenomic pipelines, One Codex [30] and metaBIT [31] (supplementary figure 6a and b) to try to reduce method-based classification biases [32]. Six samples from archival documents held in the Borthwick Archive were also included for comparison. The taxon distributions generated from the metaBIT and One Codex analysis of the York Gospel samples were found to be consistent with those previously reported for the Skin microbiome (supplementary figure 6a and b)[31,33]. This result is in agreement with the fact that the skin microbiome is the most common component of the urban microbiome [34], in particular on handled surfaces [35].

To further investigate this skin microbial signature, a principal component analysis (PCoA) of the York Gospel samples was conducted with the human microbiome project (HMP) human-associated microbial profiles provided with metaBIT at genus level (figure 2a). All the York Gospel samples irrespective of their human / endogenous DNA percentages or conservation status were found to have a microbial profile which placed them within the HMP skin and nose diversity (figure 2a). This tight clustering of the York Gospel samples is not seen in the six comparative documents, whose broader distribution may reflect their more diverse life histories prior to and within the archives (supplementary figure 7). The skin microbial signature is further highlighted in an abundance heatmap (figure 2b) which shows the presence of skin microbial markers (e.g. *Propionibacterium*, and *Staphylococcus*) at high relative frequencies. Importantly these skin microbiome signatures are absent in the control sample (supplementary figure 8) suggesting they reflect microbiota colonising the parchment itself and are not the result of laboratory contamination [36,37]. Of importance to the continued conservation of the York Gospels is the discovery of the *Saccharopolyspora* genus on all bifolio including the conserved sample folio 158. This bacterial genus has been identified previously by Piñar *et al*. [13] as a possible cause for a measles-like (maculae) spotting of parchment, which is associated with localised collagen damage and document degradation [13].

To explore shared patterns of microbial colonisation within the York Gospels, samples were clustered according to their genus profiles (dendrogram figure 2b). This analysis placed the microbial composition of the highly conserved folio 158 as an outlier to all other York Gospel samples, moreover the two later additions to the manuscript (Fol. 3 and Fol. 6) are seen to cluster. Interestingly, two samples with relatively high endogenous DNA content (Fol. 13 and 101) also fall together, a tentative hint at the possibility of future correlations between microbial colonisation and DNA retrieval. The clustering analysis was then repeated using the combined Borthwick Archive / York Gospels dataset (supplementary figure 8). In this analysis three major groupings can be seen, firstly, folio 158 is again seen as an outlier. Two further internal groupings are then revealed (supplementary figure 8); one composed of the non-conserved York Gospel samples and two Borthwick Archive samples (BA39 and BA74) and a second containing the remaining Borwick Archive samples (4/6).

To investigate the relationship between the metagenomes of the York Gospels and the Borthwick Archive samples further, STAMP [38] was used to profile the MetaPhlAn2 output. In a PCoA analysis of the samples at the genus level BA39 and BA74 were again seen to cluster with the unconserved York Gospel samples, away from the four other Borthwick Archive samples (supplementary figure 9a). When BA39 and BA74 were directly compared to the remaining Borthwick Archive samples again with STAMP (Welch's t-test) three taxa (supplementary figure 9b) were significantly differentiated between the two groups (*Granulicella*, *Saccharopolyspora* and *Pseudonocardia*). The increased prevalence of *Saccharopolyspora* in BA39 and BA74 may therefore indicate why these documents cluster with the York Gospel samples given the significant presence of this genus on the manuscript (supplementary figure 9b).

A final comparison was conducted in STAMP (Welch's t-test) between the filtered Borthwick Archive dataset (BA39 and BA74 removed) and the non-conserved York Gospel samples (supplementary figure 9c), highlighting five significantly differentiated taxa between the groups (*Saccharopolyspora*, *Pseudonocardia*, *Actinopolyspora*, *Propionibacterium*, and *Staphylococcus*). These results seem to further reflect the differences in the level of handling between the documents with the York Gospels microbiome containing significantly more of the skin microbiome components *Propionibacterium* and *Staphylococcus*, but might also provide insights into their relative conservation priorities with the York Gospels being more heavily colonised overall by *Saccharopolyspora* (supplementary figure 9c).

The limited number of significantly differentiable taxa in these metagenome comparisons may be indicative of the small sample size in this study and the inherent difficulty of metagenomic taxonomic assignment from shotgun sequencing [32]. However, it could also be an indication of a common parchment microbiome, maintained by the specific microbial growth conditions (salt rich) provided by parchments surface [13].

**Figure 2:**
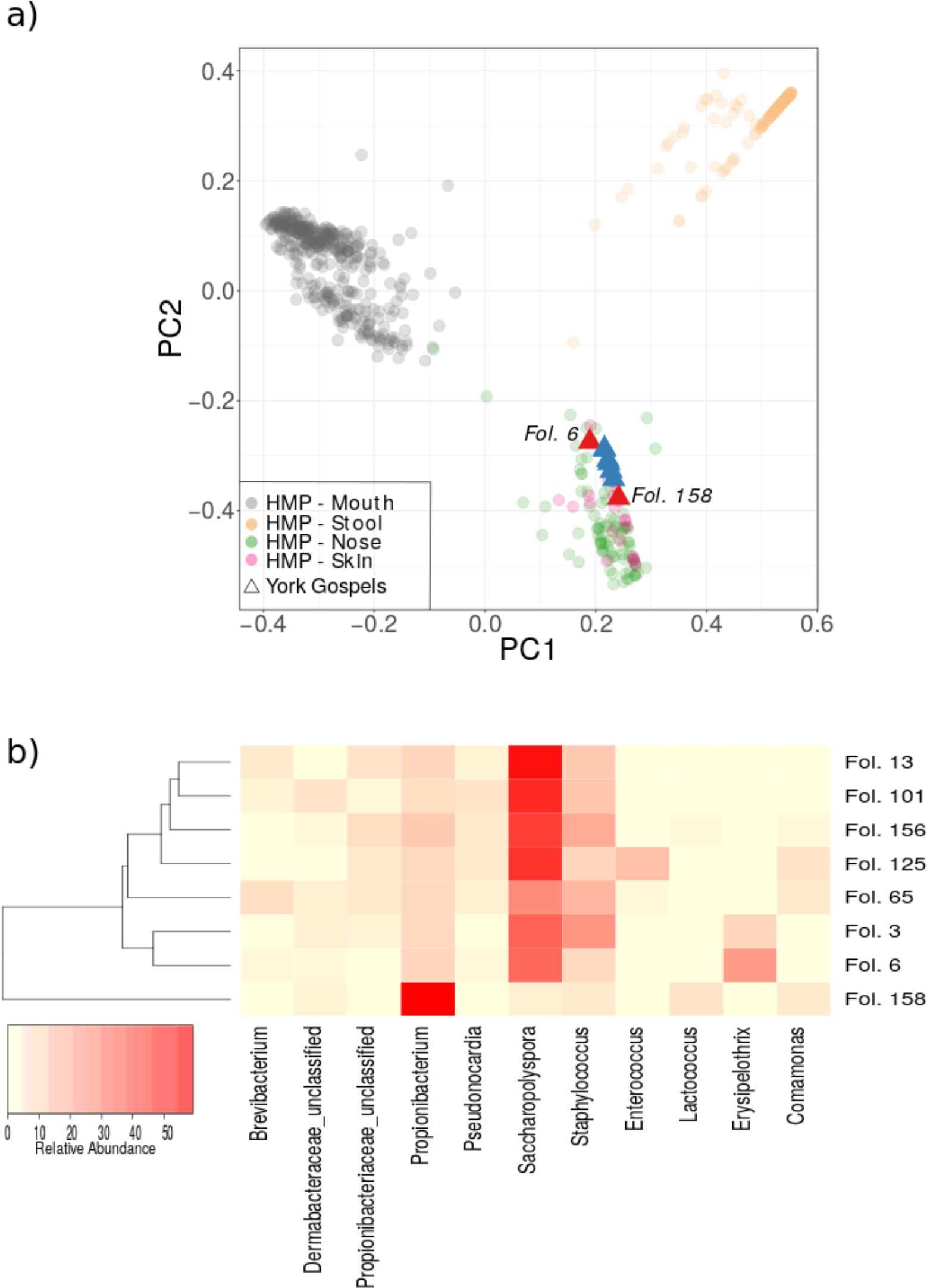
Metagenomic analysis of the York Gospel samples **a)** PCoA of York Gospels samples (host filtered) at the genus level analysed with a human microbiome project (HMP) background. The two York Gospel samples with the greatest concentration of human DNA (Fol. 6 and 158) are highlighted in red. **b)** Heatmap of microbial genus relative abundance in each of the York Gospel samples, genera shown represent >5% abundance in at least one sample. Clustering of samples (dendrogram) was completed using the complete metaBIT genus filtered output. The highly conserved sample Fol.158 is seen as an outlier to the other York Gospel samples and the two later additions (Fol. 3 and 6) are seen to cluster.

## Discussion

Illuminated manuscripts represent an irreplaceable historical record, but the need to conserve these documents is seemingly at odds with their value as an important reservoir of contemporaneous biological information. By extending the non-invasive eZooMS method to include the analysis of DNA we propose a cost effective and simple-to-use biomolecular sampling technique to enable this historical resource to be explored. The current study applies this combined methodology to document for the first time a rich palimpsest of biological information from a complete book object of great cultural value over its 1,000 year history.

### Species composition and book production

Within this study the species composition of the York Gospels has been revealed, the primary document is composed almost exclusively of calf skin, apart from a single bifolio made of sheep. This detailed analysis further highlights the utility of eZooMS for high throughput species identification of manuscripts. With a low cost and analysis time per sample, this technique has the potential to describe the species composition of many other documents and provide further insights into illuminated manuscript production.

As, zooarchaeology usually struggles to obtain accurate population sized assemblages, with collections often processed and fragmented and rarely constrained to a narrow time range [39,40]. The analysis of the species composition of manuscripts may have implications, which extend beyond the documents production, by providing a more refined understanding of past animal population sizes with a tighter chronology than can often be obtained from archaeological assemblages alone.

Although a possible artefact of the small sample number, the frequency of female animals (4 females, 1 male) among the calves is worthy of mention. As cattle are slaughtered for parchment production as juveniles (supplementary materials and methods), male calves, of lesser reproductive value than females, would be hypothesized to be most often selected for this purpose. If this contradictory pattern of an excess of female calves were to be confirmed by further sampling, a possible explanation could lie in the correlation between the writing of the manuscript and a historical outbreak of murrain, tentatively identified as rinderpest or a closely related morbillivirus ancestor. Although the composition date of the York Gospels is still debated, it is possible they were written in Canterbury around 990 CE, shortly after a major outbreak of cattle plague occurred in the Great Britain and Ireland. At least seven independent sources describe widespread cattle mortalities in England, Wales, Ireland and possibly Scotland between 986 and 988 [41,42]. Medieval and later texts demonstrate that the flaying of the carcass of a diseased animal to employ its skin was an accepted way of cutting losses. The murrain could therefore have produced an abundant source of foetal and newborn calf skins of both sexes, that were possibly so numerous they were still being used as parchment two to four years after the outbreak. Indeed, even if these skins were not produced as a consequence of the murrain but there was a widespread local mortality in the years before the text was written, it would seem unusual to sacrifice the very animals (females) required to rebuild the herd. Further analyses of the mortality patterns and local outbreaks of disease could help to refine the chronology of text.

An alternative explanation could lie in the value of the female calves themselves, as cattle are proposed to be a great source of wealth and power in the Anglo-Saxon world. If female calves are of higher value than male, perhaps the selection of female animals reflects the choice of the very best and most expensive material available to receive the holy word: a sacrifice of prized animals fit to answer the sacrifice of the Lord, and to demonstrate the faith and the wealth of the commissioner of the manuscript [43].

Finally, our assumption about the higher value of female calves, which relies mostly on data from earlier (Roman period) or later (13th-14th century) texts, may be erroneous. The Saxon economy relied mostly on oxen for traction [44], and the latter may have out-valued heifers, leading to occasional surfeits of female calves.

In this context of the perceived value of calves, the presence of a single sheepskin bifolio in the original Gospel text seems out of place. In the accounts of a Cistercian Abbey at Beaulieu (1269-1270) the best sheepskin parchment is worth less than the worst calf parchment [45]. Determining the extent to which the selection of sex and of species reflected differences in quality, perceived notions of value or a spiritual dimension would require a more comprehensive study of Anglo-Saxon manuscripts.

In relation to the later sheepskin additions to the document a 16th century inventory describes the York Gospels as “A text, decorated with silver, not well gilt, on which the oaths of the dean and other dignities and canons are inserted at the beginning” [21]. This description would seem to imply that not only the text but the bifolia themselves were later 14thC additions. A possible explanation for the use of sheepskin for these subsequent additions that contain oaths, deeds and personal correspondence is found in the *The Dialogus de Scaccario* [46], which describes a preference for legal documents to be written on sheepskin to avoid erasure and fraud of the written details. Unlike calf and goat, sheepskin has a lower density of collagen fibres at the base of the (more abundant) hair follicles, which means that the skin can split and the upper layer peel away if the parchment is roughly abraded.

### Population genetic analysis

This study is the first to our knowledge to use a non-invasive method to retrieve host genetic data from parchment, arguably the most important biological material of the Medieval world. We were able to recover endogenous DNA from all six of the samples taken from the original, almost one thousand year old York Gospel document. Although only limited DNA sequencing was undertaken in this analysis there was still sufficient data to estimate the genetic affinities of half of these samples, placing them within European cattle diversity.

The loss of more data than would be expected through standard bioinformatic filters for repetitive sequences may represent a limitation to this nucleotide retrieval technique, as the abundance of these sequences found in our dataset restricted the utility of the host genomic data. However, this increase in repetitive sequences has also been observed to a lesser extent in DNA samples from younger cut pieces of sheep parchment [1]. Given the extensive sampling opportunities that could be provided by our novel non-invasive method, a further comprehensive analyses of a diverse range of documents is warranted to see if this effect is truly a consistent artefact of eraser based extraction, or an as yet undescribed feature of DNA recovered from older parchments. Encouragingly, one sample (folio 101) did have sufficient genetic density to enable clear comparisons with reference populations and was found to closely resemble breeds of North West Europe, a result that matches the provenance of the object.

These analyses highlight a second text contained within the manuscripts of the Medieval World, which speaks to the historical management of animals and the further possibility of geolocating the source of the parchment, by comparison with the archaeological record of butchered bone. Moreover, as methods for DNA recovery and analysis are optimised further, the phenotype of animals selected for parchment production may be documented [47,48]. Genetic analyses could reveal not only the sex of the animals, but also their coat colour and morphological features (polled *vs*. non-polled), while epigenetic analyses may in the future provide insight into the animal's age [49,50].

### Metagenomics

The analysis of microbial communities that inhabit the built environment is an expanding field of research, and is moving into heritage science. We have developed an easy to use non-invasive sampling method, which can recover high resolution microbial data from sensitive documents. Our culture-free technique has benefits over other recently proposed sampling methodologies [6] in that it co-opts a widely accepted manuscript cleaning technique [20] and as such can be reliably implemented into workflows by conservators and codicologists.

One of the greatest factors influencing interpretation of microbiomes is technical variation in the analysis [51], we therefore opted for a shotgun approach to guard against known artefacts in the 16S analysis of short ancient DNA sequences [52]. Moreover, although it is known that HMWt DNA is preserved on filter paper (Owens and Szalanski 2005) (a surface in some ways analogous to parchment) we chose not to shear the extracted DNA, in an attempt to exclude recent DNA transferred by handling and preserve highly degraded endogenous molecules [53]. Reagent contamination is also a known issue for studies such as ours with limited DNA starting concentrations [36,37], which we addressed through the use of appropriate blank controls. Finally, we utilised multiple software packages in our analysis, to improve the resolution of taxon presence and abundance [32]. Despite the relative immaturity of shotgun metagenomic analyses, it is encouraging that the different analytical pipelines used in the study gave relatively consistent results, and the species of interest have overlaps with those previously described in the parchment microbiome.

The microbial signature of the York Gospel reflects the nature and use of this document in that it resembles that of the human skin microbiome, reiterated by the level of human DNA discovered on the surface of the document with values of over 15% of recovered reads in some cases. The increased abundance of the *Propionibacterium* genus seen in this analysis compared to other studies, likely reflects the significant amount of handling the York Gospels has been subject to, including its use in ecclesiastical ceremonies to this day. An alternative explanation is that the true extent of colonization by *Propionibacterium* species on parchments may have been underestimated by some 16S studies as the targeted sequencing of the hypervariable region 4 has been shown to limit the resolution of skin commensal microbiota, particularly *Propionibacterium* [54]. The discovery of the possibly destructive *Saccharopolyspora* genus [13] at high concentrations on the surface of the York Gospels further highlights the utility of metagenomic analyses that seek to describe the parchment metagenome. Illuminated manuscripts are part of our collective heritage and methods like these that can aid target conservation efforts will be of great use.

Comparison of the microbial signatures generated in this analysis highlighted a distinct difference between a highly conserved bifolio and the remaining bifolia from the York Gospels and enabled documents that were later additions to the manuscript to be distinguished. These results raise the possibility of future molecular provenancing of documents as more manuscripts have their microbiomes explored. The metagenomic signals from our comparisons of the Borthwick Archive and York Gospel samples strengthens this assertion as overall, we could distinguish between the two groups. The limited number of statically differentiated taxa between these sample sets with very divergent conservation histories raises the possibility that the microbial colonisation of documents though widespread may be limited to certain species that can tolerate the growth conditions at parchments surface [13].

## Conclusion

This study is the first non-invasive biomolecular analysis of a complete book object. Both protein and genetic analyses have been applied to reveal the animal origins of the 167 parchment folios that make up the York Gospels. This is the first time a non-invasive sampling technique has been used to recover host DNA from parchment manuscripts, which has allowed not only for the determination of source species but also an estimate of the genetic affinities and sex of the animals with limited sequencing. In addition, our novel sampling method enabled the recovery of detailed information on the microbiome associated to individual folios, which will not only be of great interest for book conservation, but also has the potential to inform about past storage and handling of book objects.

## Methods

A full description of materials and methods is provided in supplementary methods. Briefly, the York Gospels and archival documents were sampled using the dry non-invasive eraser based sampling technique of Fiddyment *et al*. [19]. With 86 folios sampled for protein analysis, and a further eight folios and six archival documents sampled for DNA analysis. eZooMS analysis of the York Gospels and archival documents was completed following the protocol of Fiddyment *et al*. [19]. DNA was extracted from the eraser crumbs following a modified version of the protocol of Fiddyment *et al*. [19] (supplementary methods). Illumina sequencing libraries were produced for each of the samples and appropriate controls following the protocol of Meyer and Kircher [55] as modified by Gamba *et al*. [56] and sequenced on an Illumina MiSeq. Raw sequencing reads were trimmed of adapter sequences using cutadapt [57] and aligned to appropriate reference genomes using BWA [58] and filtered with SAMtools [59]. DNA damage assessments were completed using mapDamage2.0 [22] and population genetic PCoA analyses with LASER2.0 [25]. Metagenomic analyses were completed from host filtered datasets (supplementary methods) using metaBIT [31], OneCodex [30] and STAMP [38].

## Acknowledgments

We thank Alison Fairburn (Borthwick Archive, University of York) for assistance with sampling of the archival documents, Mary Garrison (University of York) for helpful discussions, Sarah Griffin and Peter Young (York Minster Library) for their curatorial expertise and for facilitating sampling of the York Gospels. This work was supported by ERC Investigator grant 295729-CodeX to DGB and MJC, Marie Curie International Fellowship PALIMPSEST FP7-PEOPLE-2011-IEF 299101 and British Academy Postdoctoral Fellowship funding to SF.

## Competing interests

We have no competing interests.

## Data accessibility

Raw sequence data is deposited at the European Nucleotide Archive XXXXX, MALDI data deposited at the Dryad digital repository doi XXXX.

## Author contributions

SF produced and analysed proteomic data. MDT, VM and SF extracted DNA and made Illumina sequencing libraries. VM performed MiSeq sequencing. MDT, SF, CS, DGD and MJC analysed and or interpreted genetic data. JV performed codicological analyses. CD and SF sampled the York Gospels and JV, CCW, MC, TPN and AB provided codicological and or archaeological interpretation. MJC and DGB supervised the study. MDT, MJC, JV, CS, DGB and SF drafted the manuscript with contributions from all authors.

## References

1. Teasdale MD, van Doorn NL, Fiddyment S, Webb CC, O'Connor T, Hofreiter M, Collins MJ, Bradley DG. 2015 Paging through history: parchment as a reservoir of ancient DNA for next generation sequencing. Philos. Trans. R. Soc. Lond. B Biol. Sci. 370. (doi:10.1098/rstb.2013.0379)

2. Campana MG et al. 2010 A flock of sheep, goats and cattle: ancient DNA analysis reveals complexities of historical parchment manufacture. J. Archaeol. Sci. 37, 1317–1325. (doi:10.1016/j.jas.2009.12.036)

3. Lech T. 2016 Ancient DNA in historical parchments - identifying a procedure for extraction and amplification of genetic material. Genet. Mol. Res. 15. (doi:10.4238/gmr.15028661)

4. Burger J, Hummel S, Herrmann B. 2000 Palaeogenetics and cultural heritage. Species determination and STR-genotyping from ancient DNA in art and artefacts. Thermochim. Acta 365, 141–146. (doi:10.1016/S0040-6031(00)00621-3)

5. Manfredi M et al. 2017 A New Method for Non-Invasive Analysis of Proteins and Small Molecules from Ancient Objects. Anal. Chem. 0, null. (doi:10.1021/acs.analchem.6b03722)

6. Kraková L et al. 2017 Comparison of methods for identification of microbial communities in book collections: Culture-dependent (sequencing and MALDI-TOF MS) and culture-independent (Illumina MiSeq). Int. Biodeterior. Biodegradation (doi:10.1016/j.ibiod.2017.02.015)

7. Wandeler P, Hoeck PEA, Keller LF. 2007 Back to the future: museum specimens in population genetics. Trends Ecol. Evol. 22, 634–642. (doi:10.1016/j.tree.2007.08.017)

8. Kemp C. 2015 Museums: The endangered dead. Nature 518, 292–294. (doi:10.1038/518292a)

9. Bi K, Linderoth T, Vanderpool D, Good JM, Nielsen R, Moritz C. 2013 Unlocking the vault: next generation museum population genomics. Mol. Ecol. (doi:10.1111/mec.12516)

10. Ávila-Arcos MC et al. 2013 One hundred twenty years of koala retrovirus evolution determined from museum skins. Mol. Biol. Evol. 30, 299–304. (doi:10.1093/molbev/mss223)

11. Cassidy LM, Teasdale MD, Carolan S, Enright R, Werner R, Bradley DG, Finlay EK, Mattiangeli V. 2017 Capturing goats: documenting two hundred years of mitochondrial DNA diversity among goat populations from Britain and Ireland. Biol. Lett. 13, 20160876. (doi:10.1098/rsbl.2016.0876)

12. Sterflinger K, Pinzari F. 2012 The revenge of time: fungal deterioration of cultural heritage with particular reference to books, paper and parchment. Environ. Microbiol. 14, 559–566. (doi:10.1111/j.1462-2920.2011.02584.x)

13. Piñar G, Sterflinger K, Pinzari F. 2015 Unmasking the measles-like parchment discoloration: molecular and microanalytical approach. Environ. Microbiol. 17, 427–443.

14. Lech T. 2016 Evaluation of microbial hazard of parchment documents on an example of the 13th century Incorporation Charter for the city of Krakow. Appl. Environ. Microbiol. (doi:10.1128/AEM.03851-15)

15. Cristiani E, Borić D. 2012 8500-year-old Late Mesolithic garment embroidery from Vlasac (Serbia): Technological, use-wear and residue analyses. J. Archaeol. Sci. 39, 3450–3469.

16. Vanhaeren M, d'Errico F, van Niekerk KL, Henshilwood CS, Erasmus RM. 2013 Thinking strings: Additional evidence for personal ornament use in the Middle Stone Age at Blombos Cave, South Africa. J. Hum. Evol. 64, 500–517. (doi:10.1016/j.jhevol.2013.02.001)

17. Choyke AM, Kováts I. 2010 Tracing the personal through generations: late medieval and Ottoman combs. In Bestial Mirrors: Using Animals to construct human Identities in medieval Europe (eds AG Pluskowski, K G.K., M Kucera, M y. Bietak, I Hein), pp. 115–127. Vienna Institute for Archaeological Science.

18. Rudy KM. 2010 Dirty books : Quantifying patterns of use in medieval manuscripts using a densitometer. Journal of historians of Netherlandish Art 2, 1–26. (doi:10.5092/jhna.2010.2.1.1)

19. Fiddyment S et al. 2015 Animal origin of 13th-century uterine vellum revealed using noninvasive peptide fingerprinting. Proc. Natl. Acad. Sci. U. S. A. 112, 15066–15071. (doi:10.1073/pnas.1512264112)

20. Book and Paper Group. 1994 The Paper Conservation Catalogue. 9th edn. Washington: The American Institute for Conservation of Historic and Artistic Works. See http://www.bcin.ca/Interface/openbcin.cgi?submit=submit&Chinkey=161443.

21. Alexander J, McGurk P, Keynes S, Barr B. 1986 The York Gospels: a facsimile with introductory essays. Presentation to the members of the Roxburghe Club. See https://books.google.com/books/about/The_York_Gospels.html?hl=&id=Bp4JYAAACAAJ.

22. Jónsson H, Ginolhac A, Schubert M, Johnson PLF, Orlando L. 2013 mapDamage2.0: fast approximate Bayesian estimates of ancient DNA damage parameters. Bioinformatics 29, 1682–1684. (doi:10.1093/bioinformatics/btt193)

23. Kistler L, Ware R, Smith O, Collins M, Allaby RG. 2017 A new model for ancient DNA decay based on paleogenomic meta-analysis. Nucleic Acids Res. (doi:10.1093/nar/gkx361)

24. Sempéré G, Moazami-Goudarzi K, Eggen A, Laloë D, Gautier M, Flori L. 2015 WIDDE: a Web-Interfaced next generation database for genetic diversity exploration, with a first application in cattle. BMC Genomics 16, 940. (doi:10.1186/s12864-015-2181-1)

25. Wang C, Zhan X, Liang L, Abecasis GR, Lin X. 2015 Improved Ancestry Estimation for both Genotyping and Sequencing Data using Projection Procrustes Analysis and Genotype Imputation. Am. J. Hum. Genet. (doi:10.1016/j.ajhg.2015.04.018)

26. Li JZ et al. 2008 Worldwide human relationships inferred from genome-wide patterns of variation. Science 319, 1100–1104. (doi:10.1126/science.1153717)

27. Skoglund P, Ersmark E, Palkopoulou E, Dalén L. 2015 Ancient Wolf Genome Reveals an Early Divergence of Domestic Dog Ancestors and Admixture into High-Latitude Breeds. Curr. Biol. (doi:10.1016/j.cub.2015.04.019)

28. Hanghøj K, Seguin A, Schubert M, Madsen T, Pedersen JS, Willerslev E, Orlando L. 2016 Fast, accurate and automatic ancient nucleosome and methylation maps with epiPALEOMIX. Mol. Biol. Evol., msw184. (doi:10.1093/molbev/msw184)

29. Pedersen JS et al. 2013 Genome-wide nucleosome map and cytosine methylation levels of an ancient human genome. Genome Res. 24, 454–466. (doi:10.1101/gr.163592.113)

30. Minot SS, Krumm N, Greenfield NB. 2015 One Codex: A Sensitive and Accurate Data Platform for Genomic Microbial Identification. bioRxiv., 027607. (doi:10.1101/027607)

31. Louvel G, Der Sarkissian C, Hanghøj K, Orlando L. 2016 metaBIT, an integrative and automated metagenomic pipeline for analyzing microbial profiles from high-throughput sequencing shotgun data. Mol. Ecol. Resour. (doi:10.1111/1755-0998.12546)

32. Warinner C, Herbig A, Mann A, Fellows Yates JA, Weiß CL, Burbano HA, Orlando L, Krause J. 2016 A Robust Framework for Microbial Archaeology. Annu. Rev. Genomics Hum. Genet. (doi:10.1146/annurev-genom-091416-035526)

33. Grice EA, Segre JA. 2011 The skin microbiome. Nat. Rev. Microbiol. 9, 244–253. (doi:10.1038/nrmicro2537)

34. Stephens B. 2016 What Have We Learned about the Microbiomes of Indoor Environments? mSystems 1, e00083–16.

35. Bik HM, Maritz JM, Luong A, Shin H, Dominguez-Bello MG, Carlton JM. 2016 Microbial Community Patterns Associated with Automated Teller Machine Keypads in New York City. mSphere 1. (doi:10.1128/mSphere.00226-16)

36. Salter SJ et al. 2014 Reagent and laboratory contamination can critically impact sequence-based microbiome analyses. BMC Biol. 12, 87. (doi:10.1186/s12915-014-0087-z)

37. Laurence M, Hatzis C, Brash DE. 2014 Common contaminants in next-generation sequencing that hinder discovery of low-abundance microbes. PLoS One 9, e97876. (doi:10.1371/journal.pone.0097876)

38. Parks DH, Tyson GW, Hugenholtz P, Beiko RG. 2014 STAMP: statistical analysis of taxonomic and functional profiles. Bioinformatics 30, 3123–3124. (doi:10.1093/bioinformatics/btu494)

39. Domínguez-Rodrigo M. 2012 Critical review of the MNI (minimum number of individuals) as a zooarchaeological unit of quantification. Archaeol. Anthropol. Sci. 4, 47–59. (doi:10.1007/s12520-011-0082-z)

40. McGrory S, Svensson EM, Götherström A, Mulville J, Powell AJ, Collins MJ, O'Connor TP. 2012 A novel method for integrated age and sex determination from archaeological cattle mandibles. J. Archaeol. Sci. 39, 3324–3330. (doi:10.1016/j.jas.2012.05.021)

41. Newfield T. 2013 Early Medieval Epizootics and Landscapes of Disease: The Origins and Triggers of European Livestock Pestilences, 400-1000 CE. In Landscapes and Societies in Medieval Europe East of the Elbe: Interactions Between Environmental Settings and Cultural Transformations (eds S Kleingartner, TP Newfield, S Rossignol, D Wehner), pp. 73–113. Toronto: Pontifical Institute of Medieval Studies.

42. Newfield TP. 2015 Human–Bovine Plagues in the Early Middle Ages. J. Interdiscip. Hist. 46, 1–38. (doi:10.1162/JINH_a_00794)

43. Carver M, Garner-Lahire J, Spall C. 2016 Portmahomack on Tarbat Ness: Changing Ideologies in North-east Scotland, Sixth to Sixteenth Century AD.

44. Trow-Smith R. 1957 A history of British livestock husbandry, to 1700. Routledge & Kegan Paul Ltd.

45. Gullick M. 1991 From parchmenter to scribe. Some observations on the manufacture and preparation of medieval parchment based upon a review of the literary evidence. In Pergament. Geschichte Struktur Restaurierung Herstellungrucke (ed P Rück), pp. 145–157. Sigmaringen: Jan Thorbecke.

46. Henderson EF. 1907 Select historical documents of the Middle Ages. George Bell.

47. Fortes GG, Speller CF, Hofreiter M, King TE. 2013 Phenotypes from ancient DNA: approaches, insights and prospects. Bioessays 35, 690–695. (doi:10.1002/bies.201300036)

48. Librado P et al. 2017 Ancient genomic changes associated with domestication of the horse. Science 356, 442–445. (doi:10.1126/science.aam5298)

49. Gokhman D, Meshorer E, Carmel L. 2016 Epigenetics: It's Getting Old. Past Meets Future in Paleoepigenetics. Trends Ecol. Evol. 31, 290–300. (doi:10.1016/j.tree.2016.01.010)

50. Orlando L, Gilbert MTP, Willerslev E. 2015 Reconstructing ancient genomes and epigenomes. Nat. Rev. Genet. 16, 395–408. (doi:10.1038/nrg3935)

51. Adams RI, Bateman AC, Bik HM, Meadow JF. 2015 Microbiota of the indoor environment: a meta-analysis. Microbiome 3, 49. (doi:10.1186/s40168-015-0108-3)

52. Ziesemer KA et al. 2015 Intrinsic challenges in ancient microbiome reconstruction using 16S rRNA gene amplification. Sci. Rep. 5, 16498. (doi:10.1038/srep16498)

53. Chen L, Liu P, Evans TC, Ettwiller LM. 2017 DNA damage is a pervasive cause of sequencing errors, directly confounding variant identification. Science 355, 752–756. (doi:10.1126/science.aai8690)

54. Meisel JS, Hannigan GD, Tyldsley AS, SanMiguel AJ, Hodkinson BP, Zheng Q, Grice EA. 2016 Skin Microbiome Surveys Are Strongly Influenced by Experimental Design. J. Invest. Dermatol. 136, 947–956. (doi:10.1016/j.jid.2016.01.016)

55. Meyer M, Kircher M. 2010 Illumina Sequencing Library Preparation for Highly Multiplexed Target Capture and Sequencing. Cold Spring Harb. Protoc. 2010, db.prot5448–pdb.prot5448. (doi:10.1101/pdb.prot5448)

56. Gamba C et al. 2014 Genome flux and stasis in a five millennium transect of European prehistory. Nat. Commun. 5. (doi:10.1038/ncomms6257)

57. Martin M. 2011 Cutadapt removes adapter sequences from high-throughput sequencing reads. EMBnet.journal 17, 10–12.

58. Li H, Durbin R. 2009 Fast and accurate short read alignment with Burrows-Wheeler transform. Bioinformatics 25, 1754–1760. (doi:10.1093/bioinformatics/btp324)

59. Li H et al. 2009 The Sequence Alignment/Map format and SAMtools. Bioinformatics 25, 2078–2079. (doi:10.1093/bioinformatics/btp352)

